# Volumetric single-molecule tracking inside subcellular structures

**DOI:** 10.1101/2025.06.09.658280

**Authors:** Sam Daly, Joseph E. Chambers, Caroline Jones, Bin Fu, James D. Manton, Joseph S. Beckwith, Stefan J. Marciniak, David C. Gershlick, Steven F. Lee

## Abstract

The molecular interactions that underpin all cellular functions depend on molecular motion within three-dimensional environments. Large depth-of-field single-molecule localization microscopy (3D-SMLM) methods facilitate these measurements, but their increased optical complexity and bespoke post-processing pipelines can sacrifice important cellular context. Here we combine single-molecule light-field microscopy (SMLFM) with widefield Fourier light-field microscopy for correlative volumetric organelle imaging. The instantaneous acquisition of subcellular volumes improves the sensitivity of molecular organization and diffusion measurements through the rejection of non-specific localizations. We first demonstrate our approach by measuring the molecular organization of a nuclear-localized HaloTag protein relative to cell nuclei. Next, we characterize the molecular diffusion of the soluble protein, calreticulin, in the context of *α*_1_-antitrypsin deficiency, which revealed an increase in heterogeneous motion within endoplasmic reticulum inclusions.

## Introduction

Single-molecule localization microscopy (SMLM) enables individual biomolecules to be localized with subdiffraction resolution (1–4). Cells are inherently compartmentalized and a variety of three-dimensional (3D) SMLM strategies have emerged to capture molecular organization and diffusion away from the coverslip and within subcellular structures (5–8).

Most 3D-SMLM approaches rely on transforming the diffraction-limited pattern made by a microscope across a defined depth-of-field (DoF). This approach is known as point-spread function (PSF) engineering. Among these methods are astigmatism (~1 µm DoF) (9), the double helix PSF (~4 µm DoF) (6, 10, 11), and the tetrapod PSF (~8 µm DoF) (12, 13). Alternatively, singlemolecule light-field microscopy (SMLFM; ~8 µm DoF) uses parallax to localize molecules in 3D (14). SMLFM is based on Fourier light-field microscopy (FLFM) and enables an order-of-magnitude speed improvement at 3D-SMLM compared to more established methodologies, such as the double helix PSF (15). Therefore, SMLFM is well-suited at measuring molecular organization and diffusion in living cellular systems.

Over the years, SMLM has been used extensively to study molecular organization and dynamics close to the coverslip interface, such as the plasma membrane (17, 18), endoplasmic reticulum (ER) (19), and cytoskeleton (20, 21) because their thin and/or flat structures are amenable to the shallow DoF of widefield microscopy. However, measurements of intracellular diffusion *via* single-molecule tracking (SMT) are limited by axial motion, which causes trajectories to be truncated as molecules exit the focal plane (22). Large DoF techniques, such SMLFM, enable the quantification of molecular organization and diffusion in complex intracellular architectures, such as the Golgi apparatus. Challenges in 3D-SMLM include low contrast and the accumulation of background fluorescence from the extended imaging volume (23, 24), both of which can compromise sensitivity. In standard microscopy experiments, subcellular markers in other spectral channels improve sensitivity, but the increased optical complexity and bespoke post-processing pipelines of 3D-SMLM mean that vital spatial context is often neglected (25).

The ability to spatially segment molecular localizations in 3D would result in improved sensitivity to molecular organization and diffusion within subcellular compartments. However, optical sectioning requires axial image stacks (*i*.*e*. mechanical movement), which is slow compared to the timescales of SMLM experiments, making them prone to photobleaching-based artifacts (26). A key functional advantage of FLFM is its instantaneous acquisition of volumetric information without the need for optical sectioning (16, 27, 28). Conceptually, the challenging question of 3D spatial segmentation is addressed with a FLFM optical platform comprising two imaging channels 1) a widefield channel to instantaneously capture subcellular volumes, and 2) a channel to conduct SMLFM. A simple optical schematic is presented in Figure 1a. Volumetric reconstruction of the widefield imaging channel may be achieved with Richardson–Lucy (RL) deconvolution (16), while SMLFM data can be processed as described previously (14, 15), as shown in Figure 1b. Spatial filtration would efficiently classify localizations from other cellular locations as false positives and enhance the sensitivity of 3D-SMLM to rare processes and sub-populations. Possible research themes that would benefit from increased spatial sensitivity include the study of interactions between proteins and their binding partners (29, 30), protein trafficking along the secretory pathway (31), and subcellular membrane organization (32).

**Fig. 1.**
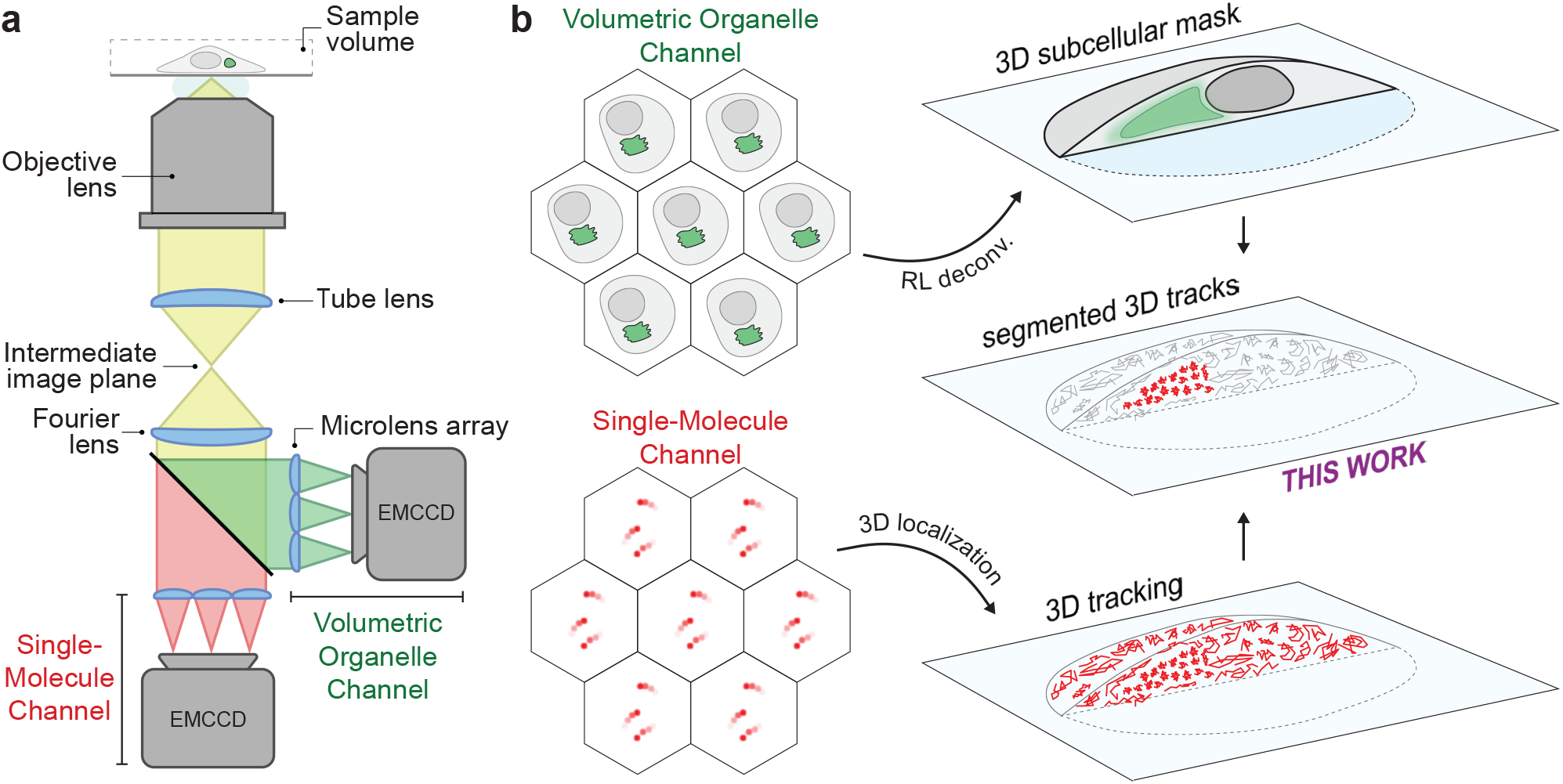
Instantaneous 3D subcellular segmentation of single-molecule fluorescence using Fourier light-field microscopy in living cells. **a** Optical schematic of two-colour Fourier light-field microscopy with key optical elements labeled. Here, one spectral channel is dedicated to widefield instantaneous volumetric imaging (Volumetric Organelle channel; green), while the other is implemented simultaneously for 3D-SMLM (Single-Molecule channel; red). **b** Analysis workflow showing 3D reconstruction of the Volumetric Organelle channel with Richardson-Lucy deconvolution (16) and the Single-Molecule channel with single-molecule 3D localization (15). Trajectories are then segmented in 3D according to their localization relative to the subcellular organelle mask for quantitative analysis with improved sensitivity.

Here, we combine SMLFM with widefield FLFM to facilitate the volumetric segmentation of localization data. First, we benchmark the resolution and reconstruction pipeline with realistic simulations and fluorescent microspheres. Next, we confirm its validity in a biological setting by measuring the spatial organization of nuclear-localized HaloTag proteins in live cell nuclei. Then we characterize the molecular motion of the soluble protein, calreticulin, in the context of α_1_-antitrypsin deficiency. Using spatial segmentation we revealed an increase in heterogeneous protein motion within endoplasmic reticulum (ER) inclusions, which supports the model of a liquid-solid phase transition (33).

## Results and Discussion

### Concept and resolution

A Fourier light-field microscopy platform was integrated into a inverted wide-field microscope. In brief, the Fourier lens was positioned in a 4f configuration with the tube lens to produce a collimated beam where a long-pass dichroic mirror was then positioned. Two identical hexagonal microlens arrays (with seven fully-illuminated lenses) were placed in the conjugate back focal plane (BFP) of both imaging channels, which focused the imaging volume onto the EMCCDs. In this work, the Volumetric Organelle (VO) channel is represented in green and the Single-Molecule (SM) channel is represented in red.

The single-molecule sensitivity and isotropic nanometer 3D resolution of the SM channel has been previously reported (15). Specifically, a resolution of ≤25 nm was demonstrated for fluorescent puncta of 3000 detected photons. This corresponds to the median signal detected from a typical single-molecule fluorophore, such as AF647, on this platform. To validate the VO channel for high-magnification volumetric imaging inside living cells we used a combination of realistic simulations and immobilized fluorescent beads. Realistic microscopy datasets were simulated as described in (15) using empirically-determined optical parameters, but with an artificially high sampling rate and brightness. This enabled the evaluation of the 3D PSF produced by RL deconvolution, see Figure 2a. The full-width half-maxima of the lateral (xy) and axial (z) components of the PSF were determined to be 406 and 597 nm, respectively. The equivalent experimental PSF (at a lower sampling rate) is presented in Figure 2b. These values quantify the smallest volumetric spatial scale that can be distinguished with the FLFM configuration. The resolution was also quantified using the Cramér-Rao lower bound as <25 nm across the whole 8 µm DoF at 4000 detected photons, as shown in Figure 2c. Finally, 4 µm fluorescent beads were imaged to confirm the accurate evaluation of particle sizes using the VO channel at the experimental sampling rate (266 nm pixel size), see Figure 2d. The superimposed circle of 4 µm diameter and accompanying 3D volume demonstrates the accurate retrieval of particle size and shape for a known geometry. Given subcellular compartments typically range in size from 0.5 µm to 10 µm, this data demonstrates that the VO channel possesses sufficient resolution to accurately capture a range of subcellular phenotypes.

**Fig. 2.**
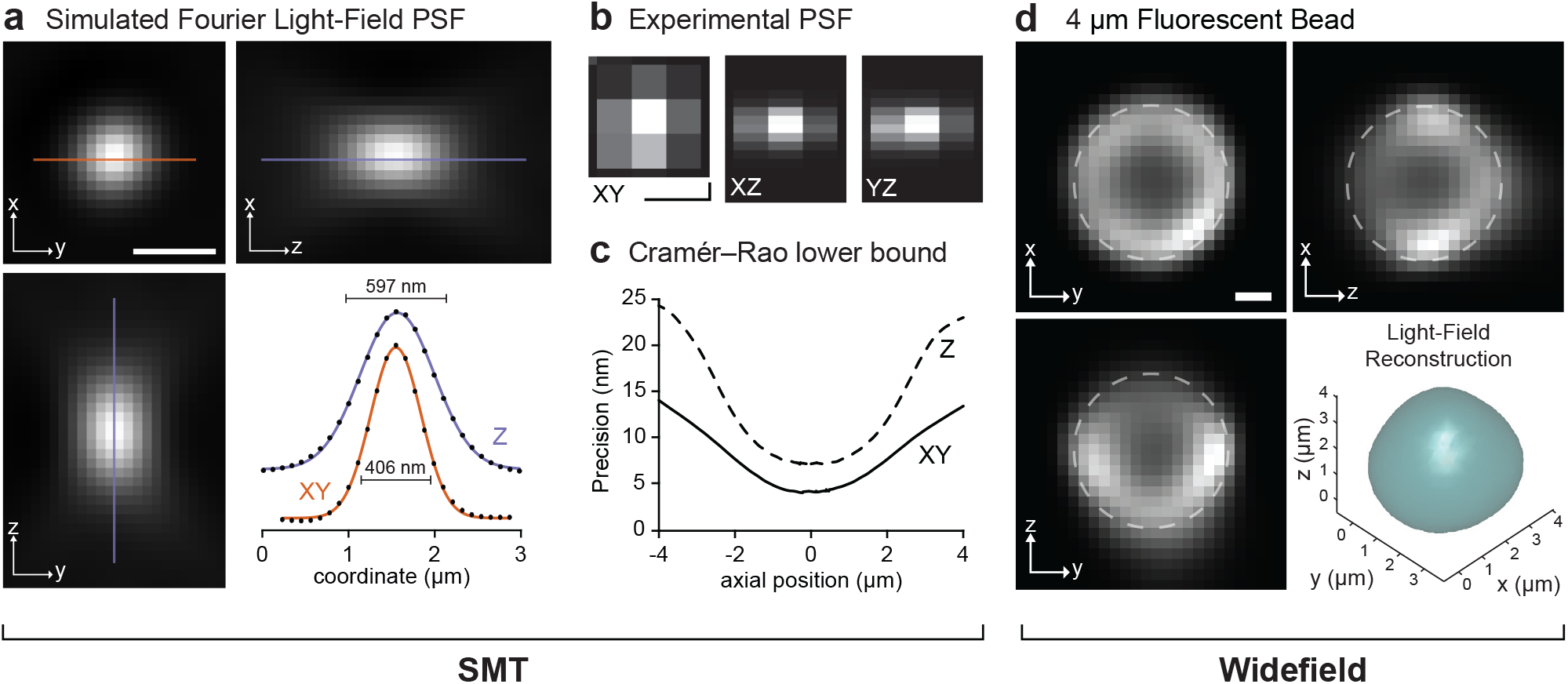
Optical validation. **a** A simulated image of the highly up-sampled 3D PSF for the optical platform described in this work. Line profiles are presented with FWHM measurements. Scale bar is 1 µm. **b** The experimental 3D PSF measured using a diffraction-limited fluorescent bead with a pixel size of 266 nm (Nyquist sampling criterion). Scale bar is 0.5 µm. **c** Simulated Cramér-Rao lower bound describing the precision obtained for a single molecule at 4000 detected photons in 3D. **d** The 3D reconstruction of a ~4 µm fluorescent bead in glycerol. Superimposed circle has a diameter of 4 µm. Scale bar is 1 µm.

### Segmentation of nuclear-localized proteins

To validate the VO channel in a biological context it was used to visualize the nuclei of live HeLa cells expressing a nuclear localization signal fused to EGFP (NLS-EGFP). Nuclear volumes were acquired at an exposure time of 20 ms and averaged over 2000 frames to improve the signal-to-noise ratio, see Figure 3a. The 3D volume measurements obtained *via* RL deconvolution closely corroborated those acquired through 40-step z-stacks using confocal microscopy, see Figure 3b & c. This validated the snapshot volumetric imaging pipeline for a biological setting, which is limited by lower SNR and more complex cellular morphologies compared to fluorescent beads.

**Fig. 3.**
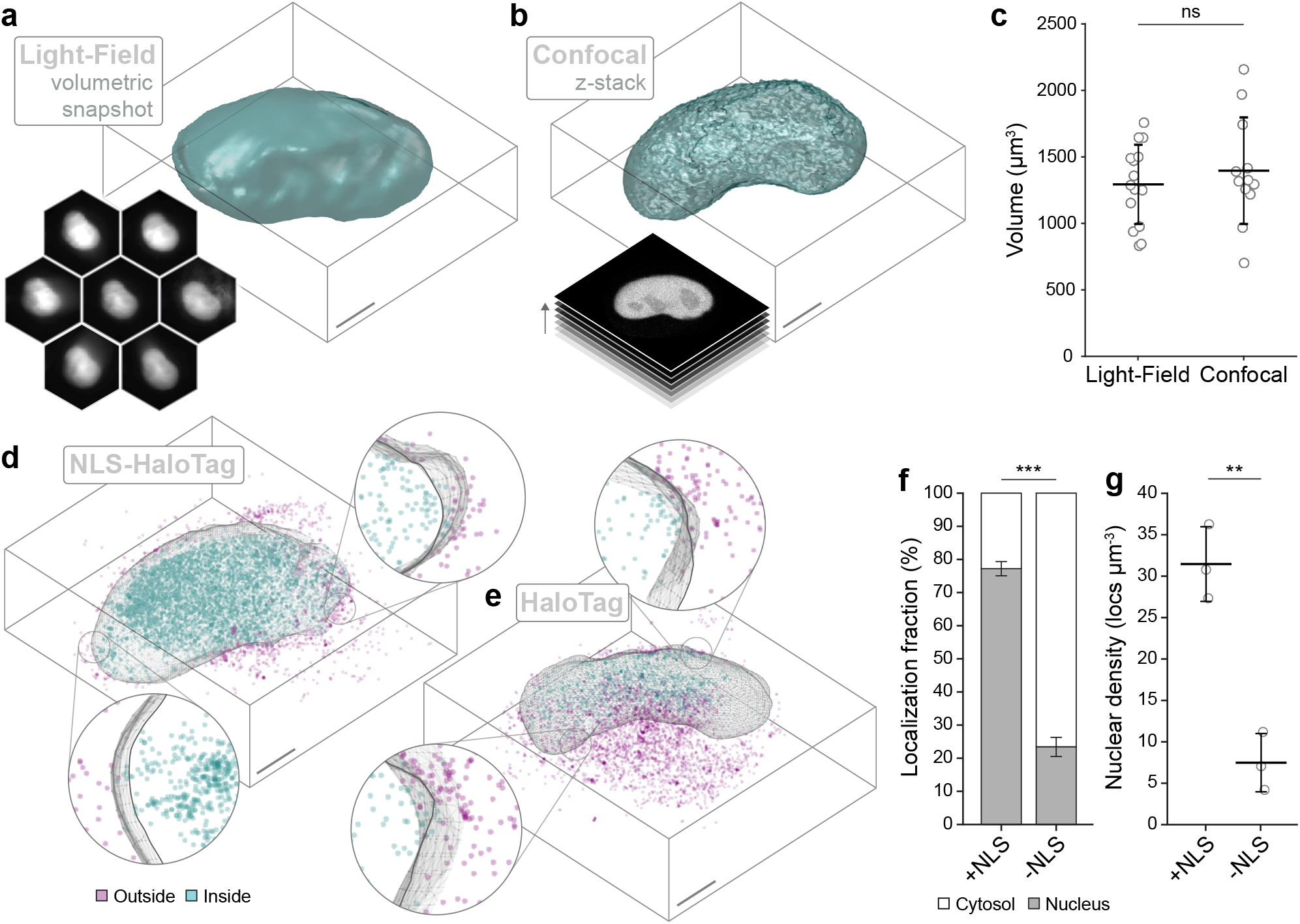
Categorizing 3D localizations according to nuclear location. **a** Representative isosurface of a HeLa cell nucleus by expression of EGFP genetically fused to a nuclear localization signal (NLS-EGFP). Volume captured instantaneously *via* Fourier light-field microscopy (FLFM, see insert, 20 µm field-of-view). **b** Representative isosurface of a HeLa cell nucleus under the same conditions *via* 43 z-planes using confocal microscopy (insert). Bounding box is ~10 µm. **c** Nuclear volumes captured by FLFM corroborate those captured by confocal z-scans. **d** Representative 3D-SMLM point cloud of NLS-HaloTag, labeled with PA-JF_646_. Localisations categorized according to appearance inside (teal) or outside (magenta) of the nuclear volume (NLS-EGFP). Insert shows cross-sections of nuclear boundary to demonstrate spatial segmentation. **e** Representative 3D-SMLM point cloud of cytosolic HaloTag (no NLS), labeled and categorized as in **d. f** Proportion of NLS-HaloTag (N = 15 cells, 3 repeats) and HaloTag localizations (N = 13 cells, 3 repeats) appearing inside and outside of the nucleus. **g** Density of NLS-HaloTag and HaloTag localizations appearing inside the nucleus relative to nuclear volume. Unless stated otherwise, data represents the average value across three biological repeats where error bars represent the standard deviation. For significance t-tests: *p < 0.05; **p < 0.01; ***p < 0.001. Scale bars are 2 µm.

Next, the VO and SM channels were implemented simultaneously to visualize cells co-expressing NLS-HaloTag and NLS-EGFP proteins, with the HaloTag labeled using PA-JF_646_ (see Methods). Laser illumination at 640, 488 and 405 nm enabled the acquisition of nuclear volumes in the VO channel and stochastic single-molecule fluorescence in the SM channel. Post-processing resulted in a 3D point cloud (NLS-HaloTag) alongside an isosurface corresponding to the nucleus (NLS-EGFP), as shown in Figure 3d. The 3D localizations were classified according to their appearance inside (teal) or outside (magenta) of the nucleus, which confirmed nuclear enrichment. The experiment was then repeated using cells co-expressing a cytosolic HaloTag protein (no NLS) alongside the NLS-EGFP. A representative 3D point cloud and nuclear isosurface is shown in Figure 3e, which now shows exclusion of localizations from the nuclear volume. The average fraction of molecules that co-localized with the nuclear volume was quantified at 77% for NLS-HaloTag, which dropped to 22% for the cytosolic HaloTag, see Figure 3f. Similarly, the density of localizations within the nucleus was 4-fold higher for NLS-HaloTag compared to the cytosolic HaloTag condition, see Figure 3g. These results indicate that the VO channel can precisely capture nuclear volumes in living cells while providing concurrent correlation with 3D-SMLM measurements.

### Measuring heterogeneous motion in the ER

Approximately 25% of all proteins transverse the secretory pathway (34). These secretory proteins fold within the endoplasmic reticulum (ER), a process that relies on the molecular motion of both the nascent polypeptide chains and ER-resident folding factors. Disruption of this process, leading to protein misfolding, is a hallmark of many human diseases (35). In α_1_-antitrypsin deficiency, the pathogenic (Z) variant aberrantly assembles into polymers within the ER of hepatocytes (36). Cells expressing Z-α_1_-antitrypsin display a range of ER morphologies, including the typical reticular network but also fragmented compartments, known as inclusions. It has been hypothesized that a protein phase transition occurs within ER inclusions, which forms a polymer matrix that restricts molecular motion (33). SMT has been used to observe the solidification of Z-α_1_-antitrypsin, but experiments were restricted to the reticular ER network due to a shallow DoF. Here, we perform volumetric SMT of the soluble, ER-localized chaperone protein, calreticulin, within individual ER inclusions for the first time and reveal heterogeneous molecular motion that is consistent with the formation of a solid protein polymer network.

The VO and SM imaging channels were implemented simultaneously to visualize living CHO-K1 cells expressing Z-α_1_-antitrypsin fused to mEmerald and calreticulin fused to HaloTag. The HaloTag protein was labeled with PA-JF_646_ to facilitate the visualization of single-molecule diffusion alongside widefield fluorescence from the ER inclusions, see Figure 4a & b (& Supplementary Movie). The mEmerald channel underwent volumetric reconstruction *via* RL deconvolution, while the single-molecule channel was super-localized and temporally grouped into trajectories, as described previously (15). The volumetric reconstruction of Z-α_1_-antitrypsin was then used to spatially segment the calreticulin trajectories according to appearance inside or outside of ER inclusions, see Figure 4c. A total of 19 cells were imaged across three biological repeats with an exposure time of 20 ms for 10,000 frames (~5 minutes) to mitigate phototoxic effects. A total of 24,836 tracks were detected, with an average length of 19.3 points per track.

**Fig. 4.**
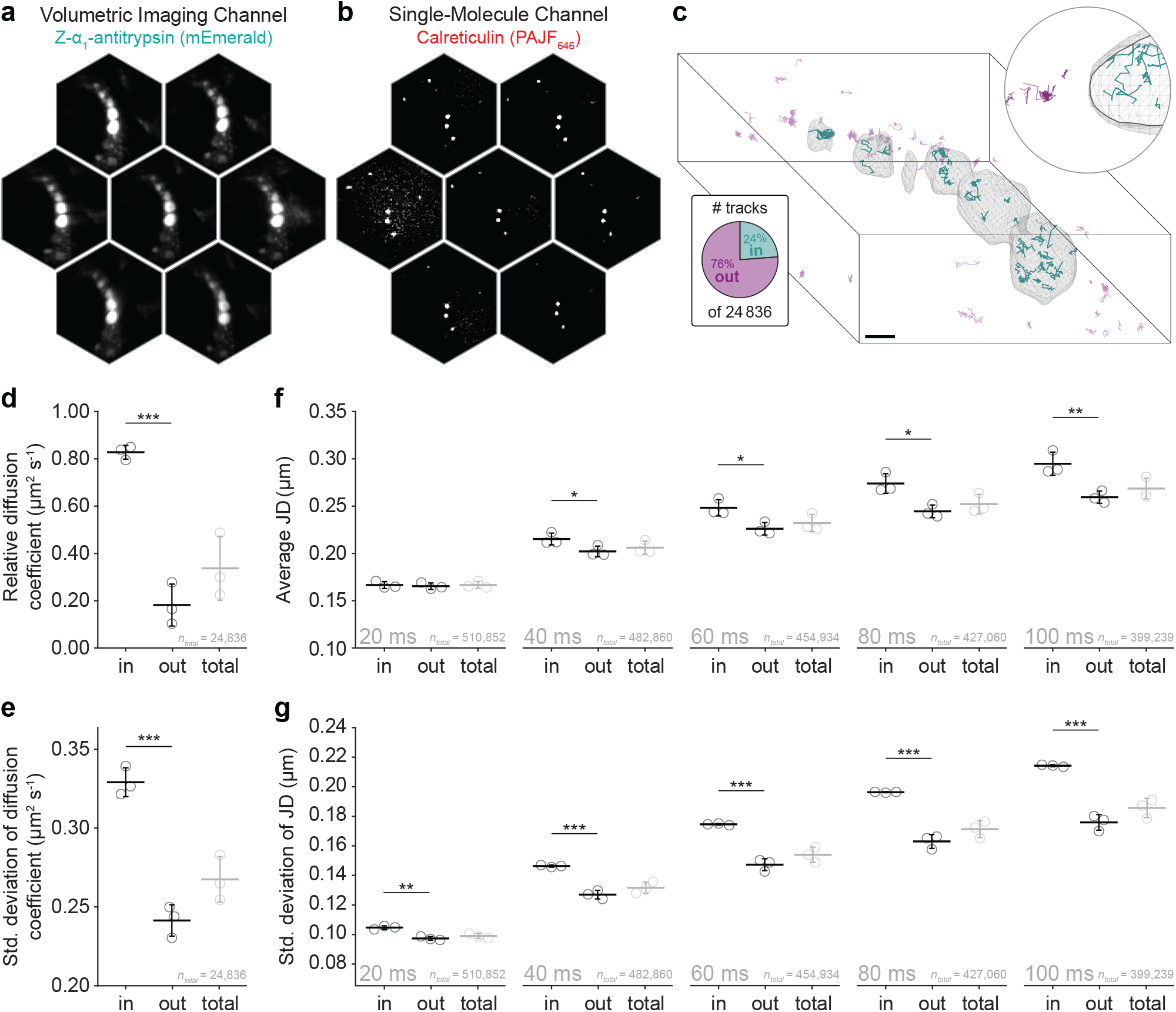
Volumetric SMT with segmentation reveals heterogeneous diffusion within the diseased ER. **a** Representative light-field image showing several ER inclusions by the expression of Z-α_1_-antitrypsin genetically fused to mEmerald. Each perspective view is 30 µm in diameter. **b** Representative single-molecule light-field image of the soluble ER protein, calreticulin, which was genetically fused to HaloTag protein and labeled with PA-JF_646_. **c** Single-molecule trajectories of calreticulin segmented by the 3D reconstruction of Z-α_1_-antitrypsin from the mEmerald channel. Teal and magenta tracks correspond to inside and outside tracks, respectively. Expanded view is 5 µm across. Representative of 2000 frames. **d** Average diffusion coefficient (mean square displacement analysis) of calreticulin inside and outside of ER inclusions, and without segmentation (total). **e** Standard deviation of diffusion coefficients in **d** within biological repeats (N = 3). **f** Average 3D jump distance inside and outside of ER inclusions, and without segmentation (total), across five lag times (Δt = 20, 40, 60, 80, and 100 ms). **g** Standard deviation of the 3D jump distances in **f** within biological repeats (N = 3) across five lag times (Δt = 20, 40, 60, 80, and 100 ms). Unless otherwise stated, error bars indicate the standard deviation across three biological repeats (totaling 19 cells or 24,836 tracks). For significance t-tests: *p < 0.05; **p < 0.01; ***p < 0.001. Sample size indicated by n_tot_. Scale bar represents 5 µm.

Mean square displacement analysis was used to evaluate the diffusion coefficient of calreticulin inside (D_in_) and outside (D_out_) of ER inclusions, see Figure 4d (& Supplementary Note 1). Diffusion within ER inclu sions, D_in_, was observed to be 1.4*×* faster than outside, D_out_. Without segmentation (D_total_), this significant difference would have been concealed within a broader distribution. The standard deviation between tracks within each biological repeat was also 1.3*×* greater for D_in_ at 0.32 µm^2^s^−1^ compared to D_out_ at 0.24 µm^2^s^−1^, as shown in Figure 4e. This broader range of diffusive behaviour is consistent with molecular motion through a cross-linked polymer matrix leading to variable confinement within ER inclusions. Importantly, no correlation was observed when diffusion coefficient was evaluated as a function of inclusion size, which lends support to the interpretation that this diffusive heterogeneity is the result of local macromolecular crowding (see Supplementary Note 2).

Jump distance (JD) analysis was then used to examine calreticulin motion on a per-step basis rather than per track, see Figure 4f (& Supplementary Note 3). Over a time lag of 20 ms (sequential frames) JD_in_ = 0.167 ± 0.004 µm and JD_out_ = 0.164 ± 0.003 µm, which suggests similar modes of molecular motion over short time scales at the resolution of the optical platform. Differences emerge as the time lag was evaluated from 20 ms to 100 ms (a five frame interval), where now JD_in_ = 0.30 ± 0.01 µm and JD_out_ = 0.26 ± 0.01 µm. This indicates that the average distance moved by calreticulin is greater inside ER inclusions over longer durations. Furthermore, the standard deviation of JDs within each sample in presented in Figure 4g. These data show that for a lag time of 20 ms, the standard deviation in JD_in_ was 0.105 µm, which doubled to 0.214 µm for a lag time of 100 ms. Meanwhile, for a lag time of 20 ms, the standard deviation in JD_out_ was 0.097 µm and 0.176 µm at 100 ms. This observed heterogeneity in jump distance supports the model in which Z-α_1_-antitrypsin undergoes a phase transition to a solid protein matrix within ER inclusions and likely reflecting differential confinement (33).

Contrast in SMLM is fundamentally constrained by the photophysical properties of fluorophores (*i*.*e*. photon budget). This challenge is amplified in SMT experiments, in which emitted photons are distributed across sequential frames, and is exaggerated further in SMLFM due to the division of fluorescence across several microlenses. An exposure time of 20 ms was selected to balance sensitivity and speed. In contrast to sCMOS detectors, the EMCCDs in this work (Evolve 512 Delta, Teledyne) operated with an EM-gain of 250 to maximize sensitivity and minimize read-out noise (see Supplementary Note 4). At 20 ms exposure, the shortest achievable exposure time was achieved that ensured a constant frame-rate. While a shorter exposure time would ensure better sampling of fast molecular motion the absolute quantitation of molecular velocity is still limited by current detector speeds and sensitivities (37). The design of brighter fluorescent probes would also facilitate faster temporal sampling. Therefore, in this work conclusions were drawn from relative heterogeneity in molecular motion rather than absolute magnitudes of speed. Importantly, without volumetric segmentation this observed heterogeneity in molecular motion is masked by the large population of calreticulin outside of ER inclusions.

In this work, we presented a simple optical method to enhance the sensitivity of 3D-SMLM to molecular organization and diffusion measurements within subcellular structures. Our method involved utilizing FLFM in a widefield mode and a single-molecule mode simultaneously, which enabled the segmentation of molecular localizations within organelles. We validated our methodology in living cellular systems through the detection of nuclear-enrichment of soluble proteins, and by measuring heterogeneous molecular motion in the diseased ER. We hope that spatially-resolved SMLFM will provide access to intracellular biomolecular organization and dynamics in a manner akin to the impact TIRF microscopy has had on studies of the plasma membrane. Future applications may span a wide range of biological disciplines, from tracking proteins at defined stages of the secretory pathway to probing organelle-specific protein–protein interactions.

## Methods

### Plasmids

Two NLS sequences (PAAKRVKLD), incorporating a flexible linker (GGSGG) and EcoRV restriction site, were inserted into the vector backbone pEGFP-N1 (Clontech) and pHALO-N1 between the EGFP/HALO and MCS genes *via* a single-step KLD reaction. pHALO-C1 and pHALO-N1 were generated by replacing the eGFP in Clontech vectors with HALO using Gibson assembly (E2621L; New England Biolabs) as described previously (38). Plasmids and primers used in this work are available upon reasonable request. All constructs were sequenced to verify their integrity.

### Culture and preparation of HeLa cells

Hela cells were cultured at 37 ° C and 5% CO_2_, in Dulbecco’s modified Eagle medium, DMEM (Gibco, 41966029), supplemented with 10% Fetal Bovine Serum (FBS; Sigma-Aldrich, F7424) and 0.2% MycoZapTM Plus-CL (Lonza, VZA–2012). Cells were passaged every three days and regularly tested for mycoplasma. For imaging, coverslips (631-0171, VWR) were cleaned under argon plasma (PDC-002, Harrick Plasma, Ithaca, NY) for 1 hour, transferred to 35 mm diameter 6-well culture dishes, and incubated with Matrigel for 1 hour (1.5 mL, 1:100 in cDMEM; Corning, 354277). Cells were seeded at a density of 0.1 *×*10^3^ cells cm-2 into fresh cDMEM and left overnight. For light-field and confocal microscopy of nuclei, cells were transfected with expression vectors encoding an NLS-EGFP fusion protein (2 µg DNA) at a 1:6 ratio of DNA to FuGene (µL). For single-molecule light-field microscopy, cells were transfected with expression vectors encoding an NLS-EGFP fusion protein (1.8 µg DNA) and a NLS-HaloTag fusion protein or HaloTag protein (0.2 µg DNA) at a 1:6 ratio of DNA to FuGene (µL, Promega). After 2 hours, the media was replaced with fresh cDMEM.

### Culture and preparation of CHO cells

Chinese Hamster Ovary (CHO) cells (Clontech) were cultured in F12 Ham’s nutrient mixture (Merck, Germany) supplemented with 10% FBS and 2 mM GlutaMAX (Thermo Fisher Scientific, USA) at 37 ° C, 5% CO_2_. For imaging, coverslips (631-0171, VWR) were cleaned under argon plasma for 1 hour and cells were seeded at a density of 1.04 *×*10^3^ cells cm^−2^ in 35 mm diameter 6-well culture dishes. 6 hours after seeding, cells were transfected with expression vectors encoding an mEmerald-A1AT fusion protein (0.5 µg DNA) and a HaloTag-Calreticulin fusion protein (0.2 µg DNA), at a 1:4 ratio of DNA (µg) to lipofectamine LTX (µl), as per the manufacturer’s instructions (Life Technology, UK). Expression vectors were reported previously in (39).

### Fluorescence labeling

Immediately before imaging, cells expressing HaloTag fusion proteins were washed with phosphate buffered saline (PBS; 1 *×* 10 mL; 14040133, ThermoFisher), labeled with PA-JF_646_ HaloTag ligand (500 nM, Janelia Materials) in OptiMEM (10149832, Gibco) for 1 minute, then washed again with PBS (5 *×* 10 mL) and returned to culture medium (1 *×* 2 mL). When required for imaging, the coverslip was transferred into an AttoFluor cell chamber (A7816, ThermoFisher), the buffer replaced with FluoroBrite (1 mL w/ 25 mM HEPES, A1896701, ThermoFisher), and transferred to a temperature controlled microscope chamber (37 ° C). Samples were imaged no longer than 2 hours after labeling.

### Preparation of fluorescent bead coverslips

Coverslips (631-1570, VWR) were cleaned under argon plasma for 1 hour, incubated with poly-L-lysine (PLL; 50 µL; P4832, Sigma-Aldrich) for 10 minutes, and were washed with PBS (3 *×* 50 µL). Fluorescent beads (0.2 µm or 4 µm; F8807 and F8859, ThermoFisher) were diluted in PBS to a concentration of 3.4 *×*10^7^ particles mL^−1^. The bead solution (50 µL) was then incubated on the PLL-coated coverslips for 3 minutes, which were then washed with PBS (3 *×* 50 µL) and transferred to the microscope for imaging.

### Single-Molecule Light-Field Microscopy optical platform

The SMLFM platform described in this work was constructed using an epi-fluorescence microscope (Eclipse Ti–U, Nikon) housing a 1.27 NA water immersion objective lens (Plan Apo VC 60*×*, Nikon, Tokyo, Japan). Excitation was achieved using 640 nm (~1 kW cm^−2^ power density, iBeam Smart-S 640-S, Toptica, Munich, Germany), 488 nm (~40 W cm^−2^ power density, Cobolt), and 405 nm (~4 W cm^−2^ power density, iBeam Smart-S 405-S, Toptica, Munich, Germany) lasers, which were circularly polarized, collimated and focused on to the back focal plane (BFP) of the objective. Unless stated otherwise, samples were excited with a highly inclined and laminated optical sheet (HILO) which was achieved by laterally displacing the excitation beam towards the edge of the BFP of the objective. The z-position of the objective was controlled with a scanning piezo (P–726 PIFOC, PI, Karlsruhe, Germany). Fluorescence was collected by the same objective and separated from the excitation beam using a quad-band dichroic mirror (Di01-R405/488/561/635-25*×*36, Semrock). The Fourier lens (*f* = 175 mm, ThorLabs) was placed in a 4f configuration with the tube lens (*f* = 200 mm, Nikon) to relay the conjugate BFP outside of the microscope body. The collimated emission beam path was then split chromatically at 90° using a long-pass dichroic (Di02–R635–25*×*36, Semrock). In the far-red emission path, a hexagonal microlens array (*f* = 175 mm, pitch = 2.39 mm) was placed in the BFP to relay the image plane onto an EMCCD (Evolve Delta 512, Photometrics, Tucson, AZ). Long-pass (BLP01-647R-25, Semrock) and band-pass (FF02-675/67-25, Semrock) emission filters were placed immediately before the detector to isolate fluorescence emission. In the short wavelength emission path, a hexagonal microlens array (*f* = 175 mm, pitch = 2.39 mm) was placed in the BFP within a rotation mount (RSP1/M, Thorlabs) to relay the image plane onto an EMCCD (Evolve Delta 512, Photometrics, Tucson, AZ) with radial alignment. Long-pass (BLP01-488R-25, Semrock) and band-pass (FF01-510/84-25, Semrock) emission filters were placed immediately before the detector to isolate fluorescence emission. The pixel size in image space was measured at 266 nm using a Ronchi ruling.

### Reconstruction of 3D-SMLM and SMT data

All experimental data were recorded as .tif stacks. 2D gaussian fitting of all emitter positions in all perspective views was carried out in Fiji using PeakFit (GDSC SMLM 2.0) to yield a set of 2D localizations for each raw frame. Given this initial set of 2D localizations, individual emitters were localized in 3D using custom MATLAB scripts available at (15) as outlined in (14). For SMT analysis, a custom-written MATLAB code was used to temporally group localisations into single trajectories (x,y,z and t), as described previously (15). Briefly, a minimum track length of 8 points was accepted using a search radius of 500 nm and allowing up to 1 consecutive dark frame. The subsequent datasets were analyzed using publicly available Python scripts at https://github.com/TheLeeLab/pyDiffusion_LeeLab. These were used to extract diffusion coefficients *via* mean square displacement analysis (40, 41) and 3D jump distances. Dark frames were accounted for through interpolation, which averaged the surrounding two tracking points to estimate the unknown position.

### Axial calibration for SMLFM

Fluorescent beads (200 nm, Deep Red FluoSpheres, ThermoFisher) were immobilized on a glass slide and imaged to calibrate for deviations in experimental and calculated the disparity from the SMLFM optical model. Glass slides were cleaned under argon plasma (PDC-002, Harrick Plasma, Ithaca, NY) for 1 hour and incubated with poly-L-lysine (PLL, 50 µL, 0.1% w/v, Sigma-Aldrich, P820) for 10 minutes. Glass slides were washed with PBS (3 *×* 50 µL) and incubated with fluorescent beads (50 µL, *ca*. 3.6 *×*10^8^ particles mL^−1^) for 3 minutes before washing further with PBS (3 *×* 50 µL). The piezo stage (P-726 PIFOC, PI, Karlsruhe, Germany) was used to scan the objective lens axially over 8 µm recording 10 frames at 30 ms exposure per 60 nm increment. The data was reconstructed in 3D and plotted against the known movement of the piezo stage. A linear fit was applied to the calibration curve, the gradient of which was a correction factor subsequently applied to all reconstructed data presented in this work.

### Richardson-Lucy deconvolution

A custom MATLAB code was developed to perform RL deconvolution. The point spread function (PSF) was simulated over a depth range from z = –5 µm to z = 5 µm, covering the whole depth of field of the imaging system. The PSF model used in deconvolution adapted the approach described in (14). To better account for spherical aberration caused by refractive index mismatch, the simulated PSF was combined with experimental data to generate a hybrid PSF (27). Voxel dimensions were set to 266 nm isotropically, matching the experimental back-projected pixel size of imaging system, and ensuring consistent spatial sampling between the image and the reconstructed volume. All reconstructions were performed on a PC with an Intel i9-10900 processor and 64 GB of RAM.

### Confocal microscopy

Volumes were acquired using a Zeiss LSM980 scanning confocal microscope with GaAsP detectors and a 63*×* 1.4 NA oil immersion objective. Samples were held at 37 ° C in cDMEM (substituted with 25 mM HEPES) during the imaging process. Images were captured using bidirectional scanning over a 512 *×* 512 frame size with a 10 ms dwell time per pixel. The back-projected pixel size was 70 nm laterally and 240 nm axially in accordance with Nyquist sampling. The 488 nm laser power density was ~2 W cm^−2^ and an average of 43 images were collected per z-stack.

### Statistics and Reproducibility

All statistical analyses were performed using a paired t-test to assess significance between conditions. Data are presented as mean *±* standard deviation unless otherwise stated. Significance thresholds of p* < 0.05, p** < 0.01, and p*** < 0.001 were used for t-tests. Sample sizes (N) are indicated in the figure legends. Experiments were repeated independently at least 3 times to ensure reproducibility.

### CODE AVAILABILITY

All relevant code and datasets used in this manuscript can be accessed *via*

Zenodo (https://doi.org/10.5281/zenodo.15602782).

## ACKNOWLEDGEMENTS

This work was supported by BBSRC awards BB/X511092/1 (SD & SFL) and UKRI715 (DCG), Royal Society grant RGF/EA/181021 (SFL), MRC awards MCMB MR/V028669/1 and MCMB MR/Y011813/1 (JEC & SJM), and the Alpha-1 Foundation (JEC & SJM; pC ID: 830153). We thank Janelia Materials for providing the PA-JF_646_ HaloTag ligand used for 3D-SPT experiments.

## AUTHOR CONTRIBUTIONS

SD, DCG and SFL conceived the project. DCG and SFL supervised the research. SD built the optical set-up, performed all experiments and analyzed data. JEC assisted in the design, implementation and analysis of tracking experiments. BF wrote the deconvolution code base. CJ assisted with the implementation of deconvolution code. JDM assisted with early deconvolution methodologies. JSB wrote and validated the pyDiffusion code base. SJM acquired funding and provided critical feedback. SD, DCG and SFL wrote the manuscript with input from all authors.

## COMPETING INTERESTS

The authors declare no conflicts of interest.

## Notes

https://doi.org/10.5281/zenodo.15602782

### Competing Interest Statement

The authors have declared no competing interest.

## References

1. Michael J Rust, Mark Bates, and Xiaowei Zhuang. Sub-diffraction-limit imaging by stochastic optical reconstruction microscopy (STORM). Nature Methods, 3(10):793–796, October 2006.

2. Eric Betzig, George H. Patterson, Rachid Sougrat, O. Wolf Lindwasser, Scott Olenych, Juan S. Bonifacino, Michael W. Davidson, Jennifer Lippincott-Schwartz, and Harald F. Hess. Imaging Intracellular Fluorescent Proteins at Nanometer Resolution. Science, 313(5793):1642–1645, September 2006. Publisher: American Association for the Advancement of Science.

3. Samuel T. Hess, Thanu P. K. Girirajan, and Michael D. Mason. Ultra-High Resolution Imaging by Fluorescence Photoactivation Localization Microscopy. Biophysical Journal, 91(11):4258–4272, December 2006.

4. Mickaël Lelek, Melina T. Gyparaki, Gerti Beliu, Florian Schueder, Juliette Griffié, Suliana Manley, Ralf Jungmann, Markus Sauer, Melike Lakadamyali, and Christophe Zimmer. Single-molecule localization microscopy. Nature Reviews Methods Primers, 1(1):39, June 2021.

5. Alex von Diezmann, Yoav Shechtman, and W. E. Moerner. Three-Dimensional Localization of Single Molecules for Super-Resolution Imaging and Single-Particle Tracking. Chemical reviews, 117(11):7244–7275, June 2017.

6. Alexander R. Carr, Aleks Ponjavic, Srinjan Basu, James McColl, Ana Mafalda Santos, Simon Davis, Ernest D. Laue, David Klenerman, and Steven F. Lee. Three-Dimensional Super-Resolution in Eukaryotic Cells Using the Double-Helix Point Spread Function. Biophysical Journal, 112(7):1444–1454, April 2017.

7. S. Basu, O. Shukron, D. Hall, P. Parutto, A. Ponjavic, D. Shah, W. Boucher, D. Lando, W. Zhang, N. Reynolds, L. H. Sober, A. Jartseva, R. Ragheb, X. Ma, J. Cramard, R. Floyd, J. Balmer, T. A. Drury, A. R. Carr, L.-M. Needham, A. Aubert, G. Communie, K. Gor, M. Steindel, L. Morey, E. Blanco, T. Bartke, L. Di Croce, I. Berger, C. Schaffitzel, S. F. Lee, T. J. Stevens, D. Klenerman, B. D. Hendrich, D. Holcman, and E. D. Laue. Live-cell three-dimensional single-molecule tracking reveals modulation of enhancer dynamics by NuRD. Nature Structural & Molecular Biology, 30(11):1628–1639, November 2023.

8. Leanne Maurice and Alberto Bilenca. Three-dimensional single particle tracking using 4π self-interference of temporally phase-shifted fluorescence. Light: Science & Applications, 12(1):58, March 2023.

9. Bo Huang, Wenqin Wang, Mark Bates, and Xiaowei Zhuang. Three-dimensional super-resolution imaging by stochastic optical reconstruction microscopy. Science (New York, N.Y.), 319(5864):810–813, February 2008.

10. Sri Rama Prasanna Pavani and Rafael Piestun. High-efficiency rotating point spread functions. Optics Express, 16(5):3484–3489, March 2008.

11. Sri Rama Prasanna Pavani, Michael A. Thompson, Julie S. Biteen, Samuel J. Lord, Na Liu, Robert J. Twieg, Rafael Piestun, and W. E. Moerner. Three-dimensional, single-molecule fluorescence imaging beyond the diffraction limit by using a double-helix point spread function. Proceedings of the National Academy of Sciences, 106(9):2995–2999, March 2009. Publisher: Proceedings of the National Academy of Sciences.

12. Yoav Shechtman, Steffen J. Sahl, Adam S. Backer, and W.E. Moerner. Optimal Point Spread Function Design for 3D Imaging. Physical Review Letters, 113(13):133902, September 2014. Publisher: American Physical Society.

13. Yoav Shechtman, Lucien E. Weiss, Adam S. Backer, Steffen J. Sahl, and W. E. Moerner. Precise Three-Dimensional Scan-Free Multiple-Particle Tracking over Large Axial Ranges with Tetrapod Point Spread Functions. Nano Letters, 15(6):4194–4199, June 2015. Publisher: American Chemical Society.

14. Ruth R. Sims, Sohaib Abdul Rehman, Martin O. Lenz, Sarah I. Benaissa, Ezra Bruggeman, Adam Clark, Edward W. Sanders, Aleks Ponjavic, Leila Muresan, Steven F. Lee, and Kevin O’Holleran. Single molecule light field microscopy. Optica, 7(9):1065–1072, September 2020. Publisher: Optica Publishing Group.

15. Sam Daly, João Ferreira Fernandes, Ezra Bruggeman, Anoushka Handa, Ruby Peters, Sarah Benaissa, Boya Zhang, Joseph S. Beckwith, Edward W. Sanders, Ruth R. Sims, David Klenerman, Simon J. Davis, Kevin O’Holleran, and Steven F. Lee. High-density volumetric super-resolution microscopy. Nature Communications, 15(1):1940, March 2024.

16. Changliang Guo, Changliang Guo, Wenhao Liu, Wenhao Liu, Xuanwen Hua, Haoyu Li, and Shu Jia. Fourier light-field microscopy. Optics Express, 27(18):25573–25594, September 2019. Publisher: Optical Society of America.

17. Christian Dietrich, Bing Yang, Takahiro Fujiwara, Akihiro Kusumi, and Ken Jacobson. Relationship of Lipid Rafts to Transient Confinement Zones Detected by Single Particle Tracking. Biophysical Journal, 82(1):274–284, January 2002.

18. Christian Niederauer, Chikim Nguyen, Miles Wang-Henderson, Johannes Stein, Sebastian Strauss, Alexander Cumberworth, Florian Stehr, Ralf Jungmann, Petra Schwille, and Kristina A. Ganzinger. Dual-color DNA-PAINT single-particle tracking enables extended studies of membrane protein interactions. Nature Communications, 14(1):4345, July 2023.

19. David Holcman, Pierre Parutto, Joseph E. Chambers, Marcus Fantham, Laurence J. Young, Stefan J. Marciniak, Clemens F. Kaminski, David Ron, and Edward Avezov. Single particle trajectories reveal active endoplasmic reticulum luminal flow. Nature Cell Biology, 20(10):1118–1125, October 2018.

20. Ahmet Yildiz, Joseph N. Forkey, Sean A. McKinney, Taekjip Ha, Yale E. Goldman, and Paul R. Selvin. Myosin V Walks Hand-Over-Hand: Single Fluorophore Imaging with 1.5-nm Localization. Science, 300(5628):2061–2065, June 2003.

21. Keith J. Mickolajczyk and William O. Hancock. Kinesin Processivity Is Determined by a Kinetic Race from a Vulnerable One-Head-Bound State. Biophysical Journal, 112(12):2615–2623, June 2017.

22. Timo Kuhn, Johannes Hettich, Rubina Davtyan, and J. Christof M. Gebhardt. Single molecule tracking and analysis framework including theory-predicted parameter settings. Scientific Reports, 11(1):9465, May 2021.

23. Robin Diekmann, Maurice Kahnwald, Andreas Schoenit, Joran Deschamps, Ulf Matti, and Jonas Ries. Optimizing imaging speed and excitation intensity for single-molecule localization microscopy. Nature Methods, 17(9):909–912, September 2020. Number: 9 Publisher: Nature Publishing Group.

24. Edward W. Sanders, Alexander R. Carr, Ezra Bruggeman, Markus Körbel, Sarah I. Benaissa, Robert F. Donat, Ana M. Santos, James McColl, Kevin O’Holleran, David Klenerman, Simon J. Davis, Steven F. Lee, and Aleks Ponjavic. resPAINT: Accelerating Volumetric Super-Resolution Localisation Microscopy by Active Control of Probe Emission**. Angewandte Chemie International Edition, 61(42):e202206919, October 2022.

25. Evan P. Perillo, Yen-Liang Liu, Khang Huynh, Cong Liu, Chao-Kai Chou, Mien-Chie Hung, Hsin-Chih Yeh, and Andrew K. Dunn. Deep and high-resolution three-dimensional tracking of single particles using nonlinear and multiplexed illumination. Nature Communications, 6(1), July 2015. Publisher: Springer Science and Business Media LLC.

26. Nadia Ruthardt, Don C Lamb, and Christoph Bräuchle. Single-particle Tracking as a Quantitative Microscopy-based Approach to Unravel Cell Entry Mechanisms of Viruses and Pharmaceutical Nanoparticles. Molecular Therapy, 19(7):1199–1211, July 2011.

27. Xuanwen Hua, Wenhao Liu, and Shu Jia. High-resolution Fourier light-field microscopy for volumetric multi-color live-cell imaging. Optica, 8(5):614–620, May 2021. Publisher: Optica Publishing Group.

28. Laura Galdón, Genaro Saavedra, Jorge Garcia-Sucerquia, Jorge Garcia-Sucerquia, Manuel Martínez-Corral, and Emilio Sánchez-Ortiga. Fourier lightfield microscopy: a practical design guide. Applied Optics, 61(10):2558–2564, April 2022. Publisher: Optica Publishing Group.

29. Ignacio Izeddin, Vincent Récamier, Lana Bosanac, Ibrahim I Cissé, Lydia Boudarene, Claire Dugast-Darzacq, Florence Proux, Olivier Bénichou, Raphaël Voituriez, Olivier Bensaude, Maxime Dahan, and Xavier Darzacq. Single-molecule tracking in live cells reveals distinct target-search strategies of transcription factors in the nucleus. eLife, 3:e02230, June 2014.

30. Joshua R. Heyza, Mariia Mikhova, and Jens C. Schmidt. Live cell single-molecule imaging to study DNA repair in human cells. DNA Repair, 129:103540, September 2023.

31. Ilya Levental and Ed Lyman. Regulation of membrane protein structure and function by their lipid nano-environment. Nature Reviews Molecular Cell Biology, 24(2):107–122, February 2023. Number: 2 Publisher: Nature Publishing Group.

32. François Simon, Lucien E. Weiss, and Sven Van Teeffelen. A guide to single-particle tracking. Nature Reviews Methods Primers, 4(1):66, September 2024.

33. Joseph E. Chambers, Nikita Zubkov, Markéta Kubánková, Jonathon Nixon-Abell, Ioanna Mela, Susana Abreu, Max Schwiening, Giulia Lavarda, Ismael López-Duarte, Jennifer A. Dickens, Tomás Torres, Clemens F. Kaminski, Liam J. Holt, Edward Avezov, James A. Huntington, Peter St George-Hyslop, Marina K. Kuimova, and Stefan J. Marciniak. Z-α1-antitrypsin polymers impose molecular filtration in the endoplasmic reticulum after undergoing phase transition to a solid state. Science Advances, 8(14):eabm2094, April 2022.

34. Danièle Stalder and David C. Gershlick. Direct trafficking pathways from the Golgi apparatus to the plasma membrane. Seminars in Cell & Developmental Biology, 107:112–125, November 2020.

35. Rammohan V Rao and Dale E Bredesen. Misfolded proteins, endoplasmic reticulum stress and neurodegeneration. Current Opinion in Cell Biology, 16(6):653–662, December 2004.

36. Catherine M. Greene, Stefan J. Marciniak, Jeffrey Teckman, Ilaria Ferrarotti, Mark L. Brantly, David A. Lomas, James K. Stoller, and Noel G. McElvaney. α1-Antitrypsin deficiency. Nature Reviews Disease Primers, 2(1):16051, July 2016.

37. Chiara Schirripa Spagnolo and Stefano Luin. Impact of temporal resolution in single particle tracking analysis. Discover Nano, 19(1), May 2024. Publisher: Springer Science and Business Media LLC.

38. Conceição Pereira, Danièle Stalder, Georgina S. F. Anderson, Amber S. Shun-Shion, Jack Houghton, Robin Antrobus, Michael A. Chapman, Daniel J. Fazakerley, and David C. Gershlick. The exocyst complex is an essential component of the mammalian constitutive secretory pathway. The Journal of Cell Biology, 222(5):e202205137, May 2023.

39. Alexander Y. Maslov, Sergey Makhortov, Shixiang Sun, Johanna Heid, Xiao Dong, Moonsook Lee, and Jan Vijg. Single-molecule, quantitative detection of low-abundance somatic mutations by high-throughput sequencing. Science Advances, 8(14):eabm3259, April 2022.

40. Xavier Michalet. Mean square displacement analysis of single-particle trajectories with localization error: Brownian motion in an isotropic medium. Physical Review E, 82(4):041914, October 2010. Publisher: American Physical Society.

41. Xavier Michalet and Andrew J. Berglund. Optimal diffusion coefficient estimation in single-particle tracking. Physical Review E, 85(6):061916, June 2012. Publisher: American Physical Society.

