## Supplementary Information for "Volumetric single-molecule tracking inside subcellular structures"

### Supplementary Information:
### Volumetric single-molecule tracking inside subcellular structures

**Sam Daly[a,b], Joseph E. Chambers[b], Caroline Jones[a], Bin Fu[a], James D. Manton[c], Joseph S. Beckwith[a], Stefan J. Marciniak[b], David C. Gershlick[b,✉], and Steven F. Lee[a,✉]**

[a]Yusuf Hamied Department of Chemistry, Lensfield Road, University of Cambridge, Cambridge, CB2 1EW, UK
[b]Cambridge Institute for Medical Research, University of Cambridge, Cambridge, CB2 0XY, UK
[c]MRC Laboratory of Molecular Biology, Francis Crick Avenue, Cambridge, CB2 0QH, UK

✉

#### Supplementary Note 1: Testing of single-molecule tracking analysis code

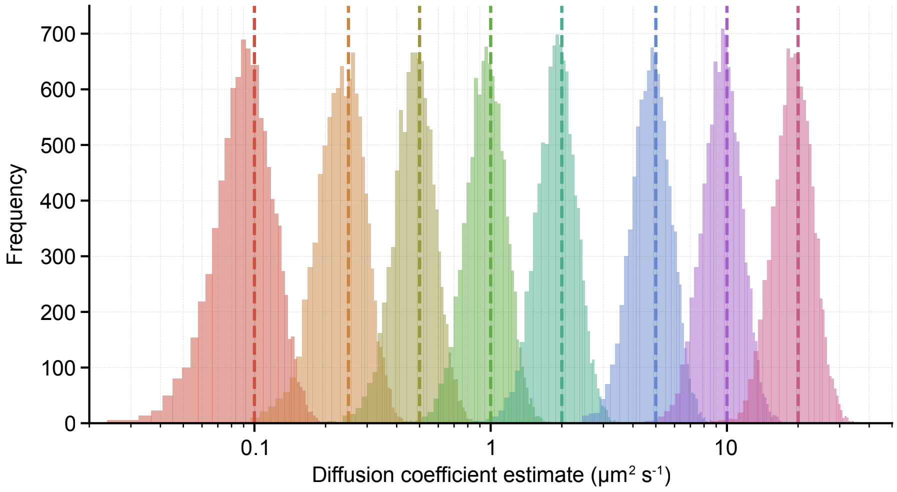


**Supplementary Figure 1. Testing of MSD analysis code.** Each color of histogram represents the estimation of a diffusion coefficient, D, from 10,000 individual trajectories (each containing 50 timesteps, displacements occurring in three dimensions) of a specified input diffusion coefficient (dotted lines). The code recapitulates the expected diffusion coefficient values. The displacements are simulated in accordance with Ref [1] assuming realistic motion blur (R=1/6, see Equation 5 of reference), a PSF with a 250 nm standard deviation and a localization precision per point of 50 nm. The timesteps of the displacements are assumed to be 1/10th of the diffusion coefficient (times micron squared) or 20 ms, whichever value was larger.

#### Supplementary Note 2: Inclusion volume does not influence the diffusion coefficient.

To confirm that segmentation does not influence the diffusion coefficient measurements in Figure 4, the inclusion volume and the diffusion coefficient of the tracks it contains should not be correlated. This relationship was evaluated and is presented in Supplementary Figure 2. Here, the diffusion coefficient was evaluated as a function of the surrounding inclusion volume, which revealed no trend. This is confirmed by a Pearson correlation coefficient of effectively zero and p-value > 0.05, see Supplementary Table 1.

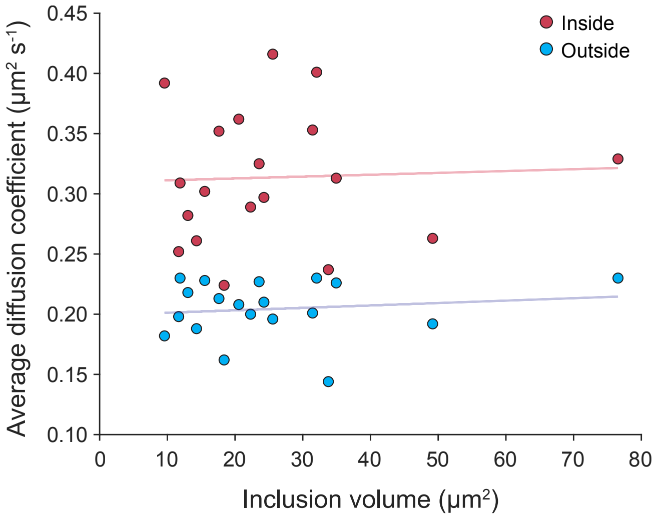


**Supplementary Figure 2. Diffusion coefficient vs. inclusion volume.** Inclusion volume was evaluated per field-of-view and is plotted here against the average diffusion coefficient of tracks. No trend is observed, which suggests that segmentation does not influence the diffusion coefficient.

**Supplementary Table 1.** Pearson correlation coefficient analysis for Supplementary Figure 2.

| | PCC | p-value |
|---|---|---|
| Outside | 0.13 | 0.59 |
| Inside | 0.04 | 0.86 |

#### Supplementary Note 3: Calreticulin diffusion is heterogeneous within ER inclusions.

The distributions of calreticulin jump distances (JD) at 20 ms and 100 ms lag times, from all experimental repeats, are shown as histograms in Supplementary Figure 3. At a lag time of $\Delta t = 20$ ms, the JD histograms for both populations are similar, suggesting comparable short-range diffusive behavior. This observation is corroborated by the overlap in their cumulative distribution functions (CDFs). Although the JDin population is smaller than JDout, the spread of JDs is comparable, indicating that both populations of motion are adequately sampled.

At longer time lag, $\Delta \mathrm{t} = 100$ms, the JDs of the two populations diverge markedly. A large immobile/highly confined fraction is observed for $JD_{out}$, with no clear change in the peak position alongside slight broadening of the distribution. These observations are consistent with the narrow, tubular geometry of the reticular ER, which restricts and adds directionality to displacement. In contrast, the distribution of $JD_{in}$ values undergoes significant broadening, leading to a large tail and small confined population. The log-scale histograms and CDFs illustrate the differences well, with $JD_{in}$ being right-shifted compared to $JD_{out}$, and a quantile difference at the 95th percentile of 0.19. These could suggest that calreticulin adopts a range of diffusive behaviors inside ER inclusions, reflecting a heterogeneous environment with variable macromolecular crowding and hence mobility. This is consistent with previous reports of molecular filtration in the ER after undergoing phase transition to a solid state [2]. Bulk fluorescence studies (*i.e.* FRAP) have been applied previously to study diffusion inside ER inclusions because single-molecule studies have been limited by the intrinsic 3D nature of these subcellular architectures that lie beyond the TIRF field.

#### Supplementary Note 4: Notes on camera gain calibrations

Conversion gain describes the relationship between counts observed in raw imaging data and the number of photons incident on the detector. The conversion gain for the EMCCDs used in this work was quantified with publicly available code (`https://github.com/TheLeeLab/cameraCalibrationCMOS`) using the method described in [3].

Full sensor images ($512 \times 512$ pixels) were recorded for 2000 frames at six exposure times (10, 20, 40, 80, 160, 320 ms), including dark frames, with electron-multiplying (EM) gain set to a value of 1. The detector was the re-calibrated with an EM gain of 250, which is the value utilized in all imaging experiments described in this work. The quantum efficiency of 0.95 at 680 nm was taken into account. Conversion gain was calculated *via* the following equation:

$$\text{conversion gain} = \text{gain (with EM gain set to 1)} \times \text{EM gain} \quad \textbf{(1)}$$

which gave values of 42 (VO channel) and 40 (SM channel) counts/photoelectron. At an EM gain of 250, average read noise was observed to be ~1.2 electrons. The in-built 'Rapid-Cal' feature of the Evolve 512 Delta ensured that the total 'conversion' gain for all experiments was consistent despite aging of the EMCCD register.

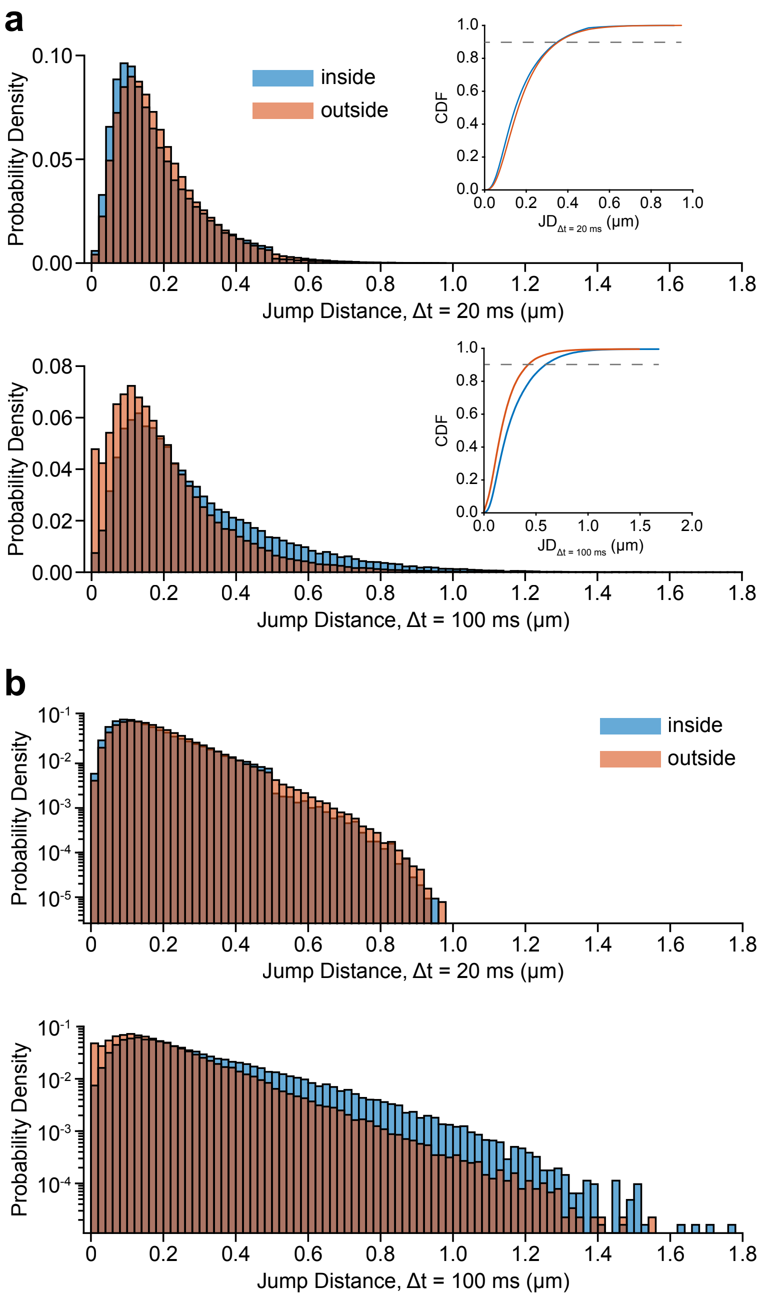


**Supplementary Figure 3. Jump distance histograms for calreticulin. a** Jump distance histogram (over a lag time of 20 ms or 100 ms) for calreticulin according to occurrence inside (blue) and outside (orange) of an ER inclusion. All tracks across N = 19 cells are shown. Insert shows the cumulative distribution function where the gray line indicates the 90th percentile. **b** Same as in **a** but on a log scale.

#### Supplementary References


1. Xavier Michalet and Andrew J. Berglund. Optimal diffusion coefficient estimation in single-particle tracking. *Physical Review E*, 85(6):061916, June 2012. Publisher: American Physical Society.
2. Joseph E. Chambers, Nikita Zubkov, Markéta Kubánková, Jonathon Nixon-Abell, Ioanna Mela, Susana Abreu, Max Schwiening, Giulia Lavarda, Ismael López-Duarte, Jennifer A. Dickens, Tomás Torres, Clemens F. Kaminski, Liam J. Holt, Edward Avezov, James A. Huntington, Peter St George-Hyslop, Marina K. Kuimova, and Stefan J. Marciniak. Z-$\alpha_1$ -antitrypsin polymers impose molecular filtration in the endoplasmic reticulum after undergoing phase transition to a solid state. *Science Advances*, 8(14):eabm2094, April 2022.
3. Fang Huang, Tobias M. P. Hartwich, Felix E. Rivera-Molina, Yu Lin, Whitney C. Duim, Jane J. Long, Pradeep D. Uchil, Jordan R. Myers, Michelle A. Baird, Walther Mothes, Michael W. Davidson, Derek Toomre, and Joerg Bewersdorf. Video-rate nanoscopy using sCMOS camera–specific single-molecule localization algorithms. *Nature Methods*, 10(7):653–658, July 2013. Number: 7 Publisher: Nature Publishing Group.
